# Cortical beta oscillations reflect the contextual gating of visual action feedback

**DOI:** 10.1101/2020.06.02.129924

**Authors:** Jakub Limanowski, Vladimir Litvak, Karl Friston

## Abstract

In sensorimotor integration, the brain needs to decide how its predictions should accommodate novel evidence by ‘gating’ sensory data depending on the current context. Here, we examined the oscillatory correlates of this process using magnetoencephalography (MEG). We used virtual reality to decouple visual (virtual) and proprioceptive (real) hand postures during a task requiring matching either modality’s grasping movements to a target oscillation. Thus, we rendered visual information either task-relevant or a (to-be-ignored) distractor. Under visuo-proprioceptive incongruence, occipital beta power decreased relative to congruence when vision was task-relevant but increased when it had to be ignored. Dynamic causal modelling (DCM) revealed that this interaction was best explained by diametrical, task-dependent changes in visual gain. These results suggest a crucial role for beta oscillations in sensorimotor integration; particularly, in the contextual gating (i.e., gain or precision control) of visual vs proprioceptive action feedback, depending on concurrent behavioral demands.

## Introduction

The integration of sensory inputs with forward or predictive models of motor control is crucial for bodily actions. In this (Bayesian) sensorimotor integration process, the brain adjusts how model predictions should respond to novel evidence by ‘gating’ sensory data, depending on the current context (Desmurget & Grafton, 2000; Friston et al., 2010;Körding & Wolpert, 2004; Sober & Sabes, 2005; Talsma et al., 2010).

Such contextual, ‘top-down’ influences on sensory gating are evinced, for instance, by the recalibration to novel (experimentally manipulated) visuo-motor mappings. Such visuo-motor recalibration has been associated with modulations of activity in visual and proprioceptive brain areas, which has been interpreted as a temporary augmentation of visual action feedback (Balslev, 2004; Bernier et al., 2009; Wasaka & Kakigi, 2012). Recently, in line with behavioral studies showing that cognitive-attentional factors can affect visuo-motor recalibration (Ingram et al., 2000; Kelso et al., 1975; Redding et al., 1985), we used fMRI to show that this activity modulation was contextual; i.e., that it depended on the relative task-relevance of seen or felt hand posture (Limanowski & Friston, 2019). However, as fMRI data provide only limited information, we could not fully characterize the fast neuronal mechanisms mediating these contextual effects.

Here, we approached this question by examining cortical oscillations with MEG. Oscillations have been linked to neuronal communication in many ways (Bressler & Richter, 2015; Buzsáki & Draguhn, 2004; de Vries et al., 2020; Donner & Siegel, 2011; Fries, 2005; Lakatos et al., 2008; Salinas & Sejnowski, 2001; Spitzer & Haegens, 2017). A widely held belief is that ‘top-down’ processes are communicated via slow oscillatory frequencies (Arnal & Giraud, 2012; Bastos et al., 2015; Donner & Siegel, 2011; Engel & Fries, 2010; Friston et al., 2015; Wang, 2010). Specifically, oscillations in the ‘alpha’ and ‘beta’ frequency ranges have been associated with sensory gating (Arnal et al., 2011; Bauer et al., 2006, 2012, 2014; Foxe et al., 1998; Foxe & Simpson, 2005; Fu et al., 2001; Haegens et al., 2012; Kelly et al., 2006; van Ede et al., 2011; Wittekindt et al., 2014) and controlling the (Kalman) gain or precision of neuronal message passing (Palmer et al., 2016; Palmer et al., 2019). Notably, changes in beta power have also been reported during movements under visuo-proprioceptive conflict (Lebar et al., 2017). Thus, oscillatory changes may indicate exactly the sort of ‘top-down’ processes that are thought to underlie the contextual gating of sensory information. However, whether this applies to the contextual (i.e., depending on cognitive-attentional factors) gating of visual action feedback during visuo-motor conflicts remains unclear.

Therefore, we recorded MEG data while participants had to match the phase of grasping movements—sensed from their unseen real hand or a seen virtual hand—to a virtual target, under varying congruence of proprioceptive (real) and visual (i.e., virtual) signals (Fig. 1A). Thus, we could examine the interaction between sensory (visuo-proprioceptive congruence) and cognitive-attentional (instructed task-relevant modality) factors. We hypothesized that—specifically under visuo-proprioceptive conflict—visual feedback should be differentially ‘gated’, depending on the current cognitive-attentional set (Corbetta & Shulman, 2002; Posner et al., 1978). Specifically, visual feedback that conflicted with the felt hand posture should be augmented when vision was task-relevant, but attenuated when vision was distracting from the task. We expected corresponding diametrical changes in induced low-frequency oscillatory power within the cortical visuo-motor hierarchy, and used DCM to disambiguate between alternative hypotheses about how these were mediated in terms of synaptic efficacy and gain control.

**Figure 1:**
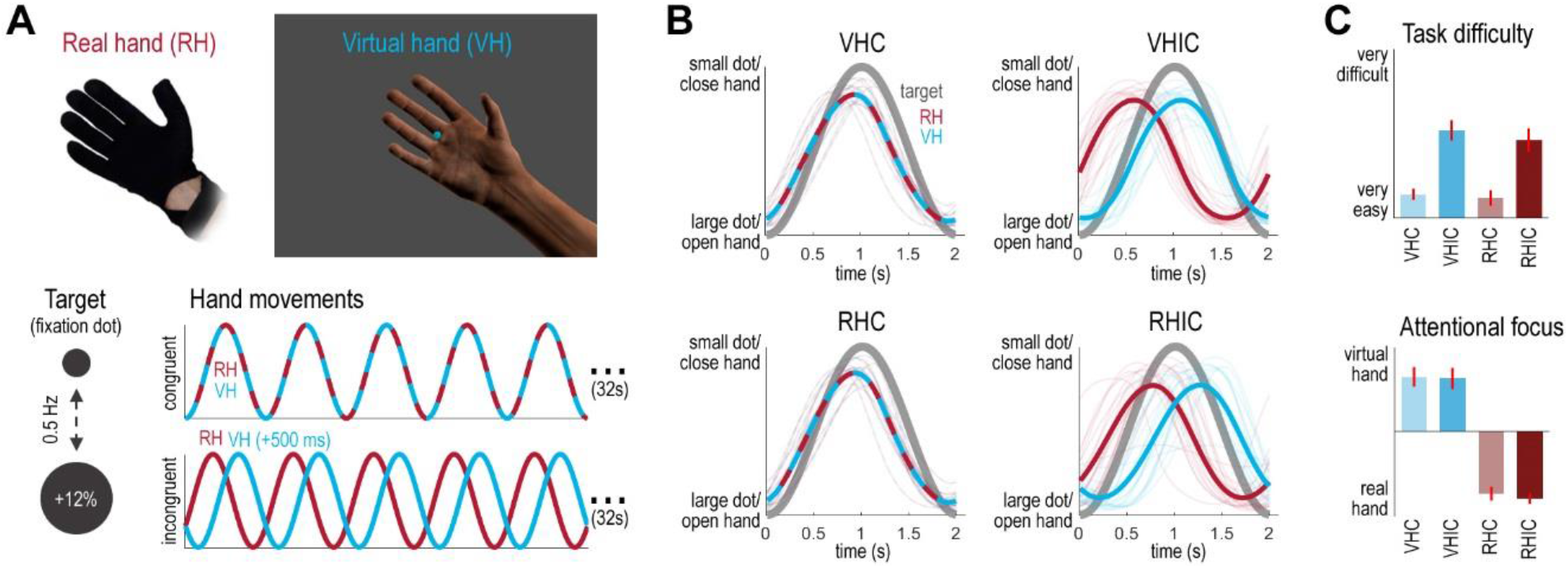
Task design and behavioral performance. **A:** Participants controlled a virtual hand (VH) model via a data glove worn on their real hand (RH), which was occluded from view. Their task was to match a 0.5 Hz oscillatory size-change of a virtual target (the fixation dot) with rhythmic grasping movements; i.e., to close the hand when the dot decreased in size and open it when the dot increased in size. Participants were instructed to align either the VH or the RH to the target oscillation in blocks of 32 s duration. In half of the conditions, the virtual hand moved congruently (C); in the other half, a 500 ms delay was added to the VH movements to introduce visuo-proprioceptive incongruence (IC). In these conditions, aligning either (virtual or real) hand with the target resulted in a misalignment of the other; consequently, participants had to select one modality over the other. We expected that under visuo-proprioceptive incongruence, visual action feedback should be differentially ‘gated’ depending on the instructed task set (VH or RH). **B:** Average movement trajectories in each condition, relative to the target’s oscillation. The individual participants’ averages are shown as thin lines. Crucially, whereas tracking was comparable when the hands moved congruently (VHC, RHC), participants exhibited phase shifts of the rhythmic movements to align the virtual hand significantly more strongly with the target in the VHIC condition, and the real hand in the RHIC condition. **C:** Participants’ mean ratings (given on 7-point visual analogue scales, with standard errors of the mean) of perceived task difficulty and attentional focus showed that visuo-proprioceptive incongruence rendered each task more difficult, and that participants complied with task instructions by directing their attention to the respective instructed hand. See Methods and Results for details.

## Materials and Methods

### Participants

18 healthy, right-handed volunteers (9 female, mean age = 29 years, range = 21-39, all with normal or corrected-to-normal vision) participated in the experiment. The sample size was determined based on a recent fMRI experiment using the same virtual reality based grasping task (Limanowski & Friston, 2019), in which we detected significant (*p* < 0.05, corrected for multiple comparisons) activations in the brain areas that were of primary interest here. The experiment was approved by the local research ethics committee (University College London) and conducted in accordance with this approval.

### Experimental design and procedure

During the experiment, participants sat underneath the MEG scanner wearing a non-magnetic data glove on their right hand, which was placed in a comfortable position on their lap and occluded from view by a black barber’s gown. The data glove (Fifth Dimension Technologies, Pretoria, South Africa; 1 sensor per finger, 8 bit flexure resolution per sensor, 60 Hz sampling rate, communication with the PC via USB with approx. 10 ms delay) measured individual finger flexions via sewn-in optical fiber cables; i.e., light was passed through the fiber cables and to one sensor per finger—the amount of light received varied with finger flexion. Prior to scanning, the glove was carefully calibrated to fit each participant’s movement range (if necessary, this was repeated between runs). The raw glove data were fed to a photorealistic virtual right hand model (Limanowski & Friston, 2019, 2020), which was thus moveable by the participant in real-time with one degree of freedom (flexion-extension) per finger. In this way, vision (*seen* hand position via the virtual hand) could be decoupled from proprioception (*felt* hand position). The virtual hand, the fixation dot, and the task instructions were presented via a projector on a screen in front of the participant (1280 × 1024 pixels resolution, screen distance to eye 64 cm, image size 40 × 29.5 cm, 32 ms projector latency). The virtual reality task environment was instantiated in the open-source 3D computer graphics software Blender (http://www.blender.org) using its Python programming interface. An eye tracker (EyeLink, SR Research) was used to monitor the participants’ eye position online, to ensure they maintained central fixation and did not close their eyes.

The participants’ task was to perform repetitive right-hand grasping movements paced by the pulsation frequency of a central fixation dot; i.e., effectively a phase matching or non-spatial pursuit task (Fig. 1A). During the movement blocks, the fixation dot continually decreased-and-increased in size sinusoidally (12 % size change) with 0.5 Hz frequency. The participants had to follow the size fluctuations with right-hand grasping movements; i.e., to close the hand when the dot shrunk and to open the hand when the dot grew. Choosing the fixation dot as the target required participants to look at the center of the screen—i.e, also at the virtual hand—under both instructions (see below), and constituted a ‘non-spatial’ target compared to e.g. targets moving along a trajectory (Limanowski et al., 2017).

In half of the movement blocks, a visuo-proprioceptive incongruence was introduced between the participant’s movements and the movements of the virtual hand; i.e., the virtual hand’s movements were delayed with respect to the actual movement by adding a 500 ms lag. In other words, the seen and felt hand positions were always incongruent (phase-shifted) in these conditions. The delay was adopted following a recent behavioral study using the similar task (Limanowski & Friston, 2020), which showed that participants reliably recognized the virtual and real hand movements as incongruent when applying this lag—and significant differences in behavior between conditions. Here, we likewise ensured that all participants were aware of the incongruence before scanning.

Crucially, participants had to perform the phase matching task with one of two goals in mind: In half of the movement blocks, they had to match the target’s oscillatory phase with the virtual hand movements or with their unseen real hand movements, respectively. This resulted in a 2 × 2 factorial design with the factors *Task (virtual hand vs real hand task)* and *Congruence (congruent vs incongruent VH/RH positions)*.

Each of the four conditions ‘virtual hand task under congruence’ (VHC), ‘virtual hand task under incongruence’ (VHIC), ‘real hand task under congruence’ (RHC), and ‘real hand task under incongruence’ (RHIC) was completed in blocks of 32 s (16 movements each) 3 times per run, in randomized order, interspersed with 6 s fixation-only periods. Participants completed five runs in total, thus completing 240 movements of 2 s each per condition. The task instructions (‘VIRTUAL’ / ‘REAL’) were presented 2.5 s before each respective movement trials for 2 seconds. Additionally, participants were informed whether in the upcoming trial the virtual hand’s movements would be synchronous (‘synch.’) or delayed (‘delay’). The instructions and the fixation dot in each task were colored (pink or turquoise, the color mapping was counterbalanced across participants), to help participants remember the current task instruction during each movement trial. Participants were trained extensively prior to scanning.

With these instructions, we aimed to induce a specific cognitive-attentional set in our participants; and with it, a different weighting of visual (vs proprioceptive) movement cues. Specifically, we hypothesized that—under visuo-proprioceptive conflict—visual action feedback should be prioritized in the VH vs RH task; i.e., depending on the currently active ‘top-down’ cognitive-attentional set (Corbetta & Shulman, 2002; Posner et al., 1978). Note that whereas in the congruent conditions, both hand positions were identical, and therefore both hands’ grasping movements could simultaneously be matched to the target’s oscillatory phase (i.e., the fixation dot’s size change), only one of the hands’ (virtual or real) movements could be phase-matched to the target in the incongruent condition. This necessarily engendered a phase mismatch of the other hand’s movements: In the VHIC condition, participants had to adjust their movements to counteract the visual lag; i.e., they had to phase-match the virtual hand’s movements (i.e., vision) to the target by shifting their real hand’s movements (i.e., proprioception) out of phase with the target. Conversely, in the RHIC condition, participants had to match their real hand’s movements (i.e., proprioception) to the target’s oscillation, and therefore had to ignore the fact that the virtual hand (i.e., vision) was out of phase. We hypothesized that this incongruence would increase task difficulty and require a sustained focus of attention on the instructed tracking modality—vision or proprioception—vs the non-instructed (‘distractor’) modality. In other words, visual feedback should be prioritized in the VHIC task (where it had to be used to recalibrate motor control to a new visuo-proprioceptive mapping) but attenuated in the RHIC task (where it was effectively distracting). In brief, we expected an interaction effect between sensory (congruence) and cognitive-attentional (task) factors.

After the experiment, participants were asked to indicate—for each of the four conditions separately—their answers to the following two questions: “How difficult did you find the task to perform in the following conditions?” (Q1, answered on a 7-point visual analogue scale from “very easy” to “very difficult”) and “On which hand did you focus your attention while performing the task?” (Q2, answered on a 7-point visual analogue scale from “I focused on my real hand” to “I focused on the virtual hand”).

### Behavioral data analysis

To analyze the behavioral change in terms of deviation from the target (i.e., phase shift from the oscillatory size change), we calculated the phase shift as the average angular difference between the raw averaged movements of the virtual or real hand (averaged over the four fingers) and the target’s oscillatory pulsation phase in each condition, using a continuous wavelet transform. The first target oscillation cycle of each block was excluded from analysis, because participants frequently only started moving with the second one. Cycles during which the hand movement was omitted (i.e., the fingers remained either flexed or extended across the entire cycle) were also excluded. On average, this left 222 trials of 2 s duration each per condition (0.4 % omitted movements).

The resulting hand phase shifts for each participant and condition were entered into a 2 × 2 repeated measures ANOVA with the factors task (virtual hand, real hand) and congruence (congruent, incongruent) to test for statistically significant group-level differences. Note that the virtual hand-target alignment directly quantified real hand-target alignment, since a larger shift of the real hand corresponds to better alignment of the virtual hand with the target. Post-hoc t-tests (two-tailed, with Bonferroni-corrected alpha levels to account for multiple comparisons) were used to compare experimental conditions. As a control analysis, we compared average movement amplitudes (i.e., the difference between maximum extension and maximum flexion per movement cycle) between conditions following the same procedure.

The questionnaire ratings were evaluated for statistically significant differences using a nonparametric Friedman’s test and Wilcoxon’s signed-rank test (with Bonferroni-corrected alpha levels to account for multiple comparisons) due to non-normal distribution of the residuals. Furthermore, we tested whether participants were inconsistent in their fixation; i.e., we tested for between-condition differences in average Euclidean distance of measured fixation from the fixation dot, using a repeated-measures 2 × 2 ANOVA analogous to the above.

### MEG data preprocessing and analysis

We hypothesized that cognitive-attentional modulations of sensory processing in our task should be reflected by changes in induced oscillatory power within the cortical visuo-motor hierarchy; specifically, in primary and extrastriate visual cortices input (Bauer et al., 2006; Foxe et al., 1998; Foxe & Simpson, 2005; Haegens et al., 2012; Lebar et al., 2017; Limanowski & Friston, 2019), temporoparietal cortex (Farrer et al., 2008; Dirk Leube et al., 2003; Limanowski et al., 2018; van Pelt et al., 2016), and the frontoparietal and prefrontal cortices (de Vries et al., 2020; Desmurget et al., 1999; Fink et al., 1999; Grefkes et al., 2004; Helfrich & Knight, 2016; Limanowski et al., 2017; Ogawa et al., 2006). To test these assumptions, we acquired MEG data during the experiment.

MEG signals were acquired using a 275-channel whole-head setup with third-order gradiometers (CTF Omega, CTF MEG International Services LP, Coquitlam, Canada) at a sampling rate of 600 Hz. All analyses were performed using MATLAB (MathWorks, Natick, MA, United States) and SPM12.6 (Wellcome Trust Centre for Neuroimaging, University College London, https://www.fil.ion.ucl.ac.uk/spm/, (Litvak et al., 2011)). MEG data were high-pass filtered (1 Hz), downsampled to 300 Hz, and epoched into trials of 2 s each (each corresponding to a full target oscillation/grasping cycle). Epochs with z-score amplitudes +-6 SD of all trials in any of the channels (8.3 % on average) were automatically rejected (Auksztulewicz et al., 2017).

In the first (in sensor space) MEG data analysis, we looked for spectral power differences between experimental conditions under ‘steady-state’ assumptions; i.e., treating the spectral profile as a ‘snapshot’ of responses induced during condition-specific changes in quasi-stationary power spectra (Donner & Siegel, 2011; Friston et al., 2019; Moran et al., 2008). We computed induced power spectra in the 0-98 Hz range using a multi-taper spectral decomposition (Thomson, 1982) with a spectral resolution of +-2 Hz. The spectra were averaged across trials using robust averaging (Litvak et al., 2012), log-transformed, and then converted to volumetric scalp × frequency images—with two spatial and one frequency dimension (Kilner & Friston, 2010). The resulting images were smoothed with a Gaussian kernel with full width at half maximum of 8 mm × 8 mm × 4 Hz and entered into a group-level general linear model (GLM) using a flexible factorial design. The statistical parametric maps obtained from the respective group-level contrasts were used to test for significant effects, using a threshold of *p* < 0.05, family-wise error (FWE) corrected for multiple comparisons at the peak level.

Following identification of regionally specific effects, source localization of induced power—in the 12-30 Hz (i.e., ‘beta’, cf. (Donner & Siegel, 2011)) range—was performed using a variational Bayesian approach with multiple sparse priors (Litvak & Friston, 2008). The 6 Hz effect for congruent > incongruent was localized separately in the 4-6 Hz range. The (source space) localization results of each participant were summarized as 3D images per condition (unsmoothed), and entered into a group-level GLM using a flexible factorial design. Since the significance of the effects on induced responses had already been established with the sensor space analysis, the source space results were displayed at a threshold of *p* < 0.005, uncorrected (*p* < 0.075 for the frontal 6 Hz-activation). The ensuing statistical parametric maps were rendered on SPM’s brain template.

### DCM of cross-spectral densities

The MEG data analysis and the analysis of our participants’ behavior suggested that—specifically under visuo-proprioceptive conflict—visual action feedback was processed (i.e., ‘gated’) differentially depending on cognitive-attentional factors (i.e., task set). This differential processing was associated with changes in oscillatory power in the ‘beta’ range over visual brain areas; i.e., there was a significant interaction effect. These results were, in principle, in line with the proposed role of low-frequency oscillations in gating sensory information flow ‘top-down’ (de Vries et al., 2020; Donner & Siegel, 2011; Engel & Fries, 2010; Friston et al., 2015; Klimesch et al., 2007; Palmer et al., 2016, 2019). To explain this effect in terms of underlying neuronal interactions, we modelled the MEG data with DCM.

DCM allows one to compare multiple alternative hypotheses (models) about how some observed data feature (in our case: spectral power across the scalp) was most likely generated by underlying interactions between and/or within neuronal populations across a network of brain sources. To model the (induced) power differences observed in the spectral analyses—in terms of source-localized neuronal interactions—we used DCM for cross-spectral densities (Friston et al., 2012; Moran et al., 2007, 2008, 2009)). This type of DCM models the synaptic mechanisms that generate spectral-domain data features and has been validated with respect to a range of previous MEG and EEG data (Auksztulewicz et al., 2017; Bastos et al., 2015; Hamburg et al., 2019; Rosch et al., 2019; Shaw et al., 2017).

We focused our DCM analysis on the crucial interaction effect identified in the spectral analysis: Beta power over occipital and temporal sensors decreased in VHIC and increased in RHIC relative to both congruent conditions. This result was in line with the behavioral results, which also showed differences between the incongruent, but not congruent, conditions. In other words, the congruent mapping conditions could be seen as a ‘baseline’ for our task, whereas the visuo-proprioceptive conflict in the incongruent conditions led to a task-dependent gating of visual information— potentially manifesting as spectral power differences—depending on the current task set. In the DCM analysis, we therefore modelled the effects of each incongruent (i.e., visuo-proprioceptive conflict) condition relative to the congruent conditions. In other words, we modelled two condition-specific effects corresponding to changes in connectivity during VHIC or RHIC, respectively, relative to the VHC and RHC conditions. To ensure optimal model fits in the frequency bands in which the spectral effects were significant (cf. Fig. 2), we modelled the 12-30 Hz range.

**Figure 2:**
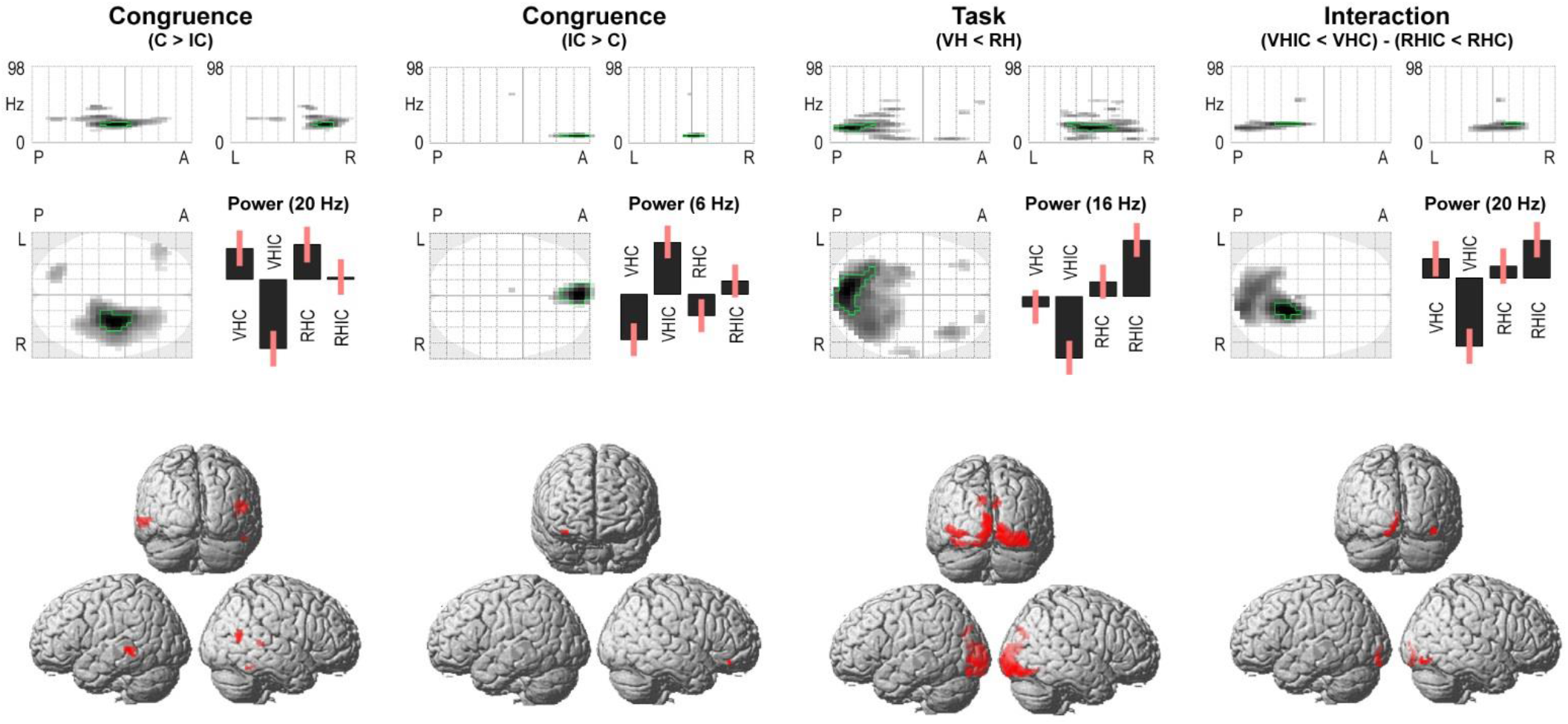
MEG spectral differences and corresponding source reconstructions. The ‘glass brain’ projections show the sensor level scalp-frequency maps of induced power differences between conditions depending on sensory (Congruence) and cognitive-attentional (Task) factors, and their interaction (maximum intensity projections displayed at *p* < 0.001, effects significant at *p*_*FWE*_ < 0.05 are outlined in green; the top plots have one frequency dimension, 0-98 Hz, and one spatial dimension, P-A = posterior-anterior, L-R = left-right; the bottom plot has two spatial dimensions). The bar plots show the mean-centered estimates of oscillatory power in each condition from the respective peak frequency, in arbitrary units and with associated standard errors. VH = virtual hand task, RH = real hand task, C = congruent hand mapping, IC = incongruent hand mapping. The renders show the corresponding source localization results using variational Laplace with multiple sparse priors. See Results for details.

Sources of interest were chosen based on the source localization of significant power differences between conditions in the above spectral analysis (see above and Fig. 2). Our DCM architecture (Fig. 3A) therefore contained the bilateral V1, V5, STS, and the right PFC; which were identified as likely sources for the main effects and, most importantly, for the interaction effect (in fact, the bilateral STS and right PFC were identified as further sources of the interaction effect, when lowering the statistical threshold of the projections to *p* < 0.1). In short, our DCM encompassed the key regions of a well-established visuo-motor hierarchy (Cisek & Kalaska, 2010; Decety et al., 1994; Goodale & Milner, 1992; Grafton, 2010; Iacoboni & Dapretto, 2006; Kilner et al., 2007; Makin et al., 2012). The reconstructed cortical locations of these effects were strikingly similar to the location of blood oxygen level dependent (BOLD) signal changes detected in our previous fMRI experiments using similar designs (Limanowski et al., 2017; Limanowski & Friston, 2019).

**Figure 3:**
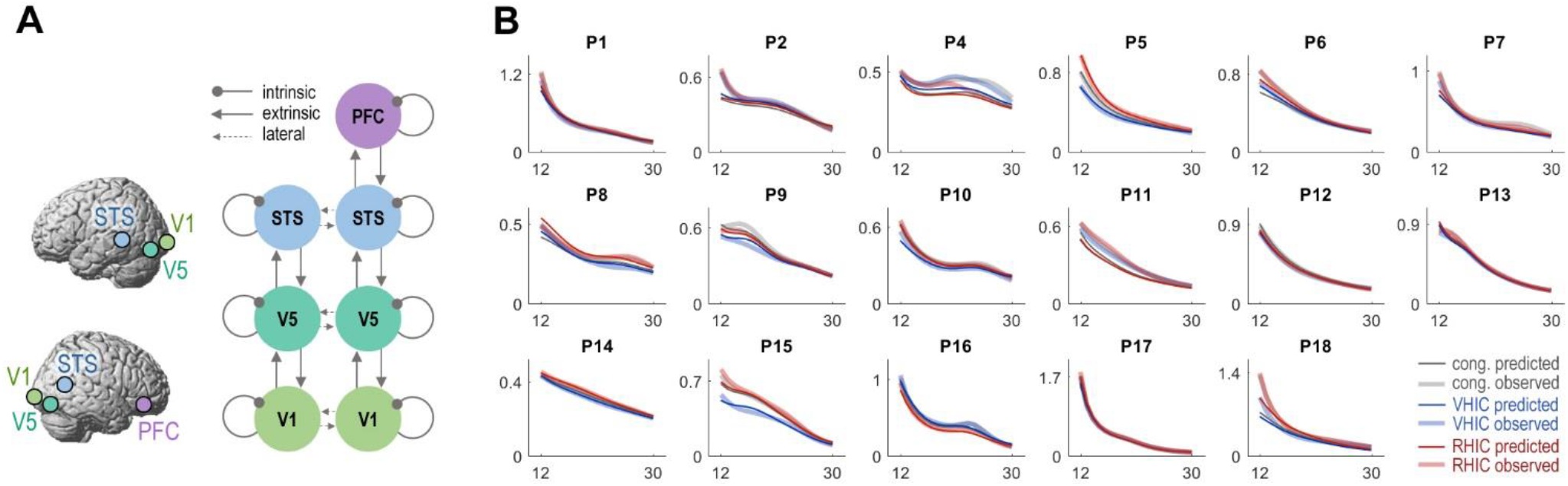
DCM architecture. **A:** A hierarchical cortical network was constructed based on the source localization of significant spectral differences between conditions; including the bilateral V1, V5, STS, and the right PFC (shown schematically, cf. Fig. 2). **B:** Individual model fits showing the first principal eigenmode of the prediction in sensor space (thin lines) and the corresponding mode of the empirical scalp data (thick lines) for the congruent ‘baseline’, and the VHIC and RHIC conditions.

Each cortical source was modelled as a local patch (Daunizeau et al., 2009; Pinotsis et al., 2012) whose responses were generated by a neural mass model (Friston et al., 2012; Moran et al., 2007) comprising three interconnected cell populations with excitatory spiny stellate cells (assigned to granular layer IV), excitatory pyramidal cells, and inhibitory interneurons (occupying both supra- and infra-granular layers). This kind of model distinguishes between ‘extrinsic’ (‘forward’ and ‘backward’) between-area connections, and ‘intrinsic’ connections. The latter connections model effects of self-inhibition, determining the input-output balance or ‘excitability’ of a given source, and are therefore usually associated with cortical gain control (Bauer et al., 2014; Pinotsis et al., 2014; Ranlund et al., 2016; Shaw et al., 2017). The resulting network allowed us to order the sources hierarchically (V1-V5-STS-PFC, with additional lateral connections for bilateral regions, Fig. 3A), and to test competing hypotheses about the type of connectivity modulation underlying the observed spectral effects.

For each participant, the resulting model (of coupled neural fields) was inverted to fit to the complex MEG cross-spectral densities in the 12-30 Hz range (as summarized by 8 principal eigenmodes) across the scalp. We excluded one participant (P3) whose data could not be fit by our model; however, this exclusion did not affect Bayesian model comparison (the same model still ‘won’ with 87% probability when P3 was included). The individual participants’ model fits—which were the basis for the Bayesian model comparison of condition-specific changes in effective connectivity—are shown in Figure 3B.

The aim of our DCM was to disambiguate between alternative explanations for how the observed effects of task instruction during incongruence on beta oscillatory power could have been mediated in terms of neuronal interactions *between* cortical regions or by changes in *local* cortical gain. Therefore, we asked whether the condition-specific effects were best explained by changes in extrinsic (forward and/or backward between-region) and/or intrinsic (within-region) connectivity (Fig. 4A). Model comparison was implemented by Bayesian model reduction (Friston et al., 2016; Friston & Penny, 2011), which allows one to compare ‘reduced’ models with variations in a subset of the ‘full’ model’s parameters. In our case, the full model allowed for modulations of all intrinsic and extrinsic connections, whereas the reduced models allowed for modulation of only one connection type, resulting in a model space of 7 models (Fig. 4A). The model with the greatest evidence (approximated via variational free-energy) was considered the ‘winning’ model. The posterior estimates of all reduced models were averaged using Bayesian model averaging (Penny et al., 2010). To confirm that the reduction in inhibition during VHIC > RHIC was significant, post-hoc t-tests were used to compare the condition-specific parameter estimates (i.e., their Bayesian model averages), under the most likely model.

**Figure 4:**
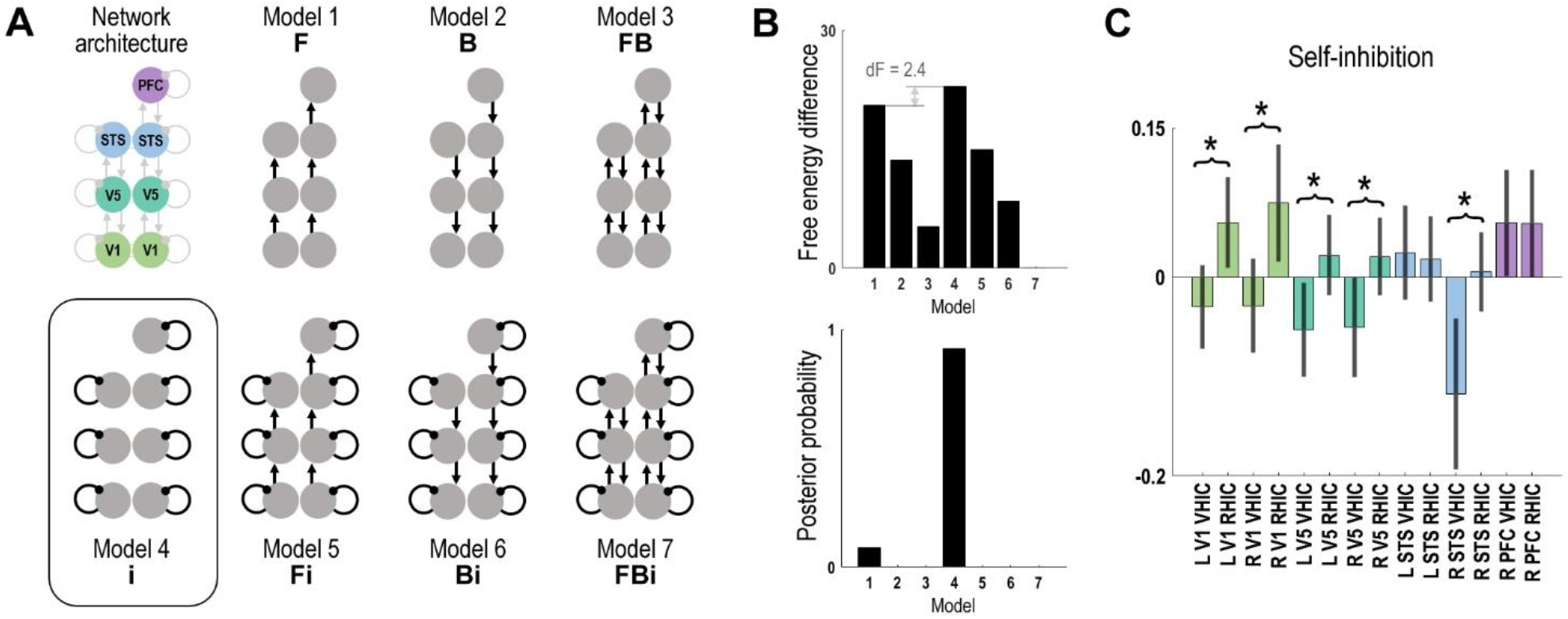
Bayesian model comparison. **A:** Using the established network architecture, 7 different candidate models were compared to test whether the condition-specific effects of VHIC and RHIC would best be modelled by changes in forward (F), backward (B), and/or intrinsic (i) connections. **B:** Bayesian model comparison identified Model 4 (condition-specific modulation of local self-inhibition) as having the highest free energy, and thus as the most likely explanation for the observed spectral data, with a posterior probability of 92%. **C:** Bayesian model averages of parameter estimates with 95% confidence intervals, indicating changes in local self-inhibition during VHIC and RHIC. Asterisks denote significantly (*p* < 0.05) reduced inhibition in VHIC relative to RHIC at the respective cortical node. See Results for details.

## Results

### Behavioral results

The participants’ average movements are shown in Figure 1B. Participants were able to track the 0.5 Hz (2 s per cycle) target oscillation with their grasping movements in all conditions, staying within ±24 degrees (~134 ms) off the target’s oscillatory phase on average. Importantly, participants aligned the virtual hand better with the target under the VH task—and, correspondingly, they aligned the real hand better with the target under the RH task (ANOVA, main effect of ‘task’, *F*_(1,16)_ = 18.11, *p* = 0.0005). A significant interaction effect between task and congruence (*F*_(1,16)_ = 18.46, *p* = 0.0005) and a post-hoc t-test showed that this effect was due to a significant difference between VHIC > RHIC (*t*_(17)_ = 4.32, *p* = 0.0004; there was no significant difference between VHC and RHC conditions, *t*_(17)_ = 0.16, *n.s.*). Unsurprisingly, tracking performance was better overall when comparing congruent to incongruent conditions (ANOVA, main effect of ‘congruence’, *F*_(1,16)_ = 136.66, *p* = 1.5e-9). Participants also evinced a partial shift of their real hand’s movements during RHIC > RHC (*t*_(17)_ = 4.43, *p* = 0.0004), but this shift was significantly smaller than during the VHIC condition (*t*_(17)_ = 4.73, *p* = 0.0002). Movement amplitudes did not differ significantly between conditions (means and standard deviations: VHC = 0.87 ± 0.05, VHIC = 0.86 ± 0.07, RHC = 0.87 ± 0.05, RHIC = 0.86 ± 0.07; repeated-measures ANOVA, all *F*_s(1,16)_ < 1, *n.s.*). Participants maintained equal fixation in all conditions (ANOVA, both main effects and interaction effect: *F*_s_ < 2.4, *n.s.*).

### The *self-reports*

(Fig. 1C) showed that, as expected, participants reported finding both tasks more difficult under visuo-proprioceptive incongruence (Friedman’s test, *χ*^2^ = 39.34, *p* < 0.001; Wilcoxon’s signed rank test, VHIC > VHC, *z*_(17)_ = 3.65, *p* < 0.001; RHIC > RHC, *z*_(17)_ = 3.31, *p* < 0.001). There was no significant difference in reported difficulty between VHC and RHC, or between VHIC and RHIC (*z*_s_ < 1.2, *n.s.*). Furthermore, as expected, participants focused their attention more strongly on the virtual hand during the virtual hand task and more strongly on the real hand during the real hand task (Friedman’s test, *χ*^2^(3,51) = 47.81, *p* < 0.001; Wilcoxon’s signed rank test, VHC > RHC, *z*_(17)_ = 3.75, *p* < 0.001) and incongruent (VHIC > RHIC, *z*_(17)_ = 3.65, *p* < 0.001) movement trials. There were no significant differences between VHC vs VHIC, and RHC vs RHIC, respectively (z_s_ < 1.2, *n.s.*).

Together, the above results suggest that, as expected, participants adopted a specific attentional set to prioritize the instructed target modality, and that this was associated with significantly better target tracking with the instructed modality (vision or proprioception) under intersensory conflict.

### MEG results

In the sensor space analysis, we looked for induced spectral power differences related to the effects of our experimental manipulations; i.e., the differential processing of incongruent vs congruent visual action feedback depending on the currently active cognitive-attentional set (VH or RH task).

This analysis revealed significant spectral correlates of visuo-proprioceptive congruence (Fig. 2): Movements under visuo-proprioceptive incongruence were associated with relatively suppressed power in the 18-22 Hz range (peak at 20 Hz) over right temporal sensors (*T* = 5.35, *p*_*FWE*_ < 0.05). These effects were source-localized to the bilateral temporal regions, focused on the superior temporal sulcus (STS). Conversely, we observed a power increase at 6 Hz (*T* = 5.79, *p*_*FWE*_ < 0.05) over frontal sensors during incongruent > congruent movements; source-localized to regions in the right prefrontal cortex (PFC). Furthermore, we found a significant effect of cognitive-attentional task set: During the VH task > RH task, power in the 12-20 Hz range (peak at 16 Hz) was significantly suppressed over occipital sensors (*T* = 5.53, *p*_*FWE*_ < 0.05, Fig. 2). These effects were source-localized to distinct peaks in the bilateral primary (V1) and extrastriate (V3, V5). The reconstructed cortical sources of beta suppression during VH > RH were almost identical to the locations of fMRI activations identified in similar fMRI tasks and contrasts (Limanowski et al., 2017; Limanowski & Friston, 2019).

Crucially, there was a significant interaction effect at 20 Hz over occipital sensors (*T* = 5.00, *p*_*FWE*_ < 0.05). This effect was localized to the left V1 and right V5. In other words, the low-frequency suppression observed during the VH task > RH task was significantly stronger during incongruent > congruent conditions. In fact, beta power suppression at temporal and occipital sensors was markedly significant in the VHIC > RHIC contrast (*T* = 6.35 and 6.96, respectively, both *p*_*FWE*_ < 0.05), but nonsignificant in the VHC > RHC contrast, even at *p* < 0.001, uncorrected. In other words, the spectral effects were largely due to a difference between the incongruent conditions—in which there was a visuo-proprioceptive conflict—with small or no differences between the congruent conditions.

### DCM results

To disambiguate between alternative hypotheses about how the interaction effect in the beta range was mediated in terms of changes in neuronal message passing among key cortical sources, we modelled the measured MEG data with DCM for cross-spectral densities. Specifically, we aimed at clarifying the nature of the task-dependent gating of visual information during VHIC and RHIC, respectively, relative to a congruent-movement ‘baseline’.

Based on the localization of the most likely sources of the observed spectral power differences, we constructed a hierarchical network comprising bilateral V1, V5, STS, and the right PFC (Fig. 3A). Using this network, the model inversion provided overall good fits of the empirical whole-scalp data in the beta range (except for participant P3 who was excluded from further analysis, see Methods). The individual model fits are shown in Figure 3B.

Based on the established model architecture, we compared alternative hypotheses about how the identified modulations of induced spectral responses during VHIC and RHIC were caused in terms of neuronal interactions. We considered a model space of 7 models (Fig. 4A), each modelling the condition-specific effects as changes in (intrinsic) synaptic efficacy *within* regions and/or (forward and/or backward) synaptic connectivity *between* regions. Model comparison, implemented using Bayesian model reduction, showed that the most likely model (Model 4, posterior probability = 92%) described the condition-specific effects in terms of a modulation of local intrinsic connections (Fig. 4B). In the DCM framework, these connections determine the degree of self-inhibition—and therefore determine input-output balance or excitability—in other words, changes of their parameter estimates indicate changes in gain control (K. Friston et al., 2015; Moran et al., 2007). The DCM results therefore suggest that, relative to a congruent-movement ‘baseline’, movements under visuo-proprioceptive conflict were most likely associated with changes in cortical gain.

Crucially, there was a striking asymmetry in the winning model’s connectivity estimates for VHIC vs RHIC (Fig. 4C): Whereas most of the intrinsic connections mediated a disinhibition during VHIC relative to the congruent movements, they showed the opposite effect—an increased self-inhibition—during RHIC. This effect was significant in the bilateral visual areas (V1 and V5) and in the right STS. In other words, relative to the baseline condition, cortical gain in visual areas was increased during VHIC and decreased during RHIC. In sum, the DCM results indicated a contextual effect of cognitive-attentional task set on cortical gain control within the visuomotor hierarchy.

## Discussion

Using a virtual reality based phase matching task under visuo-proprioceptive incongruence, we induced a cognitive-attentional prioritization of visual vs proprioceptive feedback, as evident from significant differences in target-tracking performance and self-reported attentional allocation. We found that sensory (visuo-proprioceptive congruence) and cognitive (instructed task set) factors and, importantly, their interaction effect were associated with significant changes in cortical oscillatory power—most prominently, in the ‘beta’ frequency range.

By isolating the interaction effect between sensory and cognitive-attentional factors, our study design allowed us to advance on previous work on visuo-motor recalibration: Relative to the congruent movement conditions, occipital beta power was *suppressed* in VHIC but *enhanced* in RHIC. Our DCM analysis identified diametrical changes in the cortical gain of visual areas as the most likely causes of these spectral differences; i.e., increased gain during VHIC and decreased gain during RHIC relative to movements without visuo-proprioceptive conflict. These effects were strongest in visual (V1, V5) and multisensory (right STS) areas, which are all known to process visual bodily information (Farrer et al., 2008; Lebar et al., 2017; Leube et al., 2003; Limanowski et al., 2018; Limanowski & Friston, 2019).

We, therefore, propose that these results directly reflect the contextual gating of visual bodily action information—during identical (conflicting) visuo-proprioceptive mapping—for integration with the current action plan, depending on the prevalent cognitive-attentional set. In other words, we propose that attenuated beta was associated with an increased sensitivity to visual feedback (via increased gain) when visual feedback had to be incorporated into the goal-directed action plan (VHIC), and conversely, enhanced beta was associated with an attenuation of visual feedback (via reduced neuronal gain of visual brain areas) when the visual movement was an incongruent ‘distractor’ (RHIC). Such attentional gating was not required in conditions without visuo-proprioceptive conflict (VHC and RHC).

Further support for our interpretation of beta power as being inversely related to sensory gating comes from our finding that the reconstructed cortical sources of beta *suppression* were almost identical to the locations of BOLD signal *increases* in very similar ‘vision-prioritizing’ tasks (Limanowski et al., 2017; Limanowski & Friston, 2019). This inverse relationship between source-localized beta and the BOLD signal has been reported before (Moosmann et al., 2003; Scheeringa et al., 2011; Yuan et al., 2010; Zaretskaya & Bartels, 2015) and is consistent with proposals that a loss of low-relative to high-frequency power may be associated with brain ‘activation’ detected with fMRI (Chawla et al., 1999; Kilner et al., 2005; Laufs et al., 2003; Laufs, 2008).

Thus, our results support—in a sensorimotor setting—the hypothesized link between beta oscillations and top-down contextual control in service of conveying behavioral context to lower sensory regions (Bressler & Richter, 2015; Buschman & Miller, 2007; Clark et al., 2015; Donner & Siegel, 2011; Friston et al., 2015; Spitzer & Haegens, 2017). Previous work has shown that task-irrelevant sensory brain regions can be disengaged by increasing low-frequency oscillatory activity, whereas low-frequency suppression can make stimulus processing more efficient (de Vries et al., 2020; Frey et al., 2015; Jensen & Mazaheri, 2010; Klimesch et al., 2007; Schubert et al., 2009). Although such effects are frequently observed at ‘alpha’ frequencies, beta oscillations have also been linked to active suppression of sensory input that is deemed distractive (de Vries et al., 2018; Engel & Fries, 2010; Kelly et al., 2006). In visual paradigms, attention to target stimuli while ignoring distractors suppressed occipital beta band power (Fries, 2001). In multisensory tasks, several studies have reported a negative association between parieto-occipital alpha/beta power and attention to visual—as opposed to auditory or tactile—input (Bauer et al., 2006, 2012; Foxe et al., 1998; Foxe & Simpson, 2005; Fu et al., 2001; Haegens et al., 2012; Wittekindt et al., 2014). A recent study found suppressed occipital beta power during movements under visuo-proprioceptive conflict (Lebar et al., 2017). Our findings now show that beta oscillations can be directly linked to the ‘top-down’ gating of (conflicting) visual action feedback depending on concurrent behavioral context.

Furthermore, our results support computational models following the ‘predictive coding’ framework that link slow vs fast frequency oscillations to asymmetrical message passing of predictions and errors, respectively (Arnal & Giraud, 2012; Bastos et al., 2012; Friston, 2008; Lee et al., 2013; Wang, 2010). Finally, our results speak to the particular role of beta synchronization in mediating the precision of message passing during motor control (Palmer et al., 2019). However, note that the desynchronization phenomena in our paradigm originated in the visual, as opposed to the motor system; the absence of motor beta oscillatory effects may be due to our studies’ focus on induced power during repetitive movements. Future work using complementary task designs will have to evaluate whether sensory and motor beta oscillations may in fact serve different functional roles, as has been speculated (Kilavik et al., 2013; Palmer et al., 2019; Press et al., 2011; Tan et al., 2016).

Furthermore, complementary task designs should also be used to clarify whether the (prefrontal) low-theta power increase under visuo-proprioceptive incongruence could potentially be related to the prioritization of different target information, as in visual working memory tasks (Daitch et al., 2013; Johnson et al., 2017; Liesefeld et al., 2014; Riddle et al., 2020; Sauseng et al., 2010). Finally, it should be noted that participants exhibited a partial shift of their grasping phase in the RHIC condition, which could indicate a difficulty to fully ignore matching biological motion (Borroni et al., 2005; Kilner et al., 2007, 2003). Although this should be pursued, it does not pose a problem here because behavior still differed significantly between tasks.

In conclusion, our findings suggest a critical role for beta oscillations in sensorimotor integration; i.e., indicating the ‘gating’ of visual (vs proprioceptive) action feedback depending on the current behavioral demands.

## Acknowledgements

We would like to thank Felix Blankenburg for providing the data glove.

## Author contributions

JL & KF designed study; JL acquired and analyzed data; JL wrote manuscript; VL & KF commented on the manuscript.

